# Biochemical paedomorphosis and genetic assimilation in the hypoxia adaptation of Tibetan antelope

**DOI:** 10.1101/2020.03.20.000075

**Authors:** Anthony V. Signore, Jay F. Storz

## Abstract

Developmental shifts in stage-specific gene expression can provide a ready mechanism of phenotypic change by altering the rate or timing of ontogenetic events. We discovered that the high-altitude Tibetan antelope (*Panthelops hodgsonii*) has evolved an adaptive increase in blood–O_2_ affinity by truncating the ancestral ontogeny of globin gene expression such that a high-affinity juvenile hemoglobin isoform (isoHb) completely supplants the lower-affinity isoHb that is expressed in the adult red blood cells of other bovids. This juvenilization of blood properties represents a canalization of an acclimatization response to hypoxia that has been well-documented in adult goats and sheep. We also discovered the genomic mechanism underlying this regulatory isoHb switch, revealing how a reversible acclimatization response became genetically assimilated as an irreversible adaptation to chronic hypoxia.

## Introduction

When members of multigene families are developmentally regulated, shifts in the stage-specific expression of individual genes can provide a ready mechanism of phenotypic change by altering the rate or timing of ontogenetic events (heterochrony). For example, retained activity of early-expressed genes in later stages of ontogeny can result in the retention of juvenile characters into adulthood, a well-documented developmental mechanism of phenotypic evolution (*1, 2*). In extreme cases, deceleration of development can produce a truncation of the ancestral ontogeny, resulting in the juvenilization of the adult-expressed phenotype, a phenomenon known as paedomorphosis.

In jawed vertebrates, the subfamilies of genes that encode the α- and β-type subunits of tetrameric hemoglobin (Hb) are developmentally regulated such that structurally and functionally distinct α_2_β_2_ Hb isoforms (isoHbs) are expressed during different ontogenetic stages. During mammalian development, different pre- and postnatally expressed isoHbs have evolved different oxygenation properties and perform distinct O_2_–scavenging/O_2_–transport tasks during different ontogenetic stages (*3*-*6*). Genetically based shifts in stage-specific isoHb expression could therefore provide a heterochronic mechanism of evolutionary change in respiratory gas transport and aerobic metabolism. Similarly, in humans, hereditary persistence of fetal Hb alleviates the severity of thalassemias and other pathologies affecting the synthesis or stability of adult Hb (7).

Given that prenatally expressed isoHbs of eutherian mammals exhibit substantially higher O_2_–affinities than adult-expressed isoHbs, and given that an increased Hb–O_2_ affinity is generally beneficial under conditions of severe hypoxia due to the importance of safeguarding arterial O_2_ saturation (*6, 8*-*11*), the retention of early isoHb expression into adulthood could provide an effective mechanism of adaptation to chronic O_2_ deprivation. Consistent with this hypothesis, when adult goats and sheep are exposed to acute hypoxia, they upregulate a juvenile isoHb at the expense of the normal adult isoHb (*12, 13*). Here we report the discovery of a canalized version of this response in the high-altitude Tibetan antelope, *Panthelops hodgsonii* (Artiodactyla: Bovidae), a champion among mammals in aerobic exercise performance under hypoxia. This species is endemic to the Tibetan Plateau and lives at altitudes 3600-5500 m above sea level. At an altitude of 5500 m, the partial pressure of O_2_ (*P*O_2_) is roughly half the value at sea level, a level of hypoxia that severely compromises aerobic exercise performance in humans and most other mammals (*14*-*15*). Remarkably, however, at these altitudes Tibetan antelope can sustain running speeds of >70 km/h over distances of >100 km (*16*).

In addition to documenting the phenotypic consequences of developmentally displacing the low-affinity adult isoHb with a higher-affinity juvenile isoHb – a form of biochemical paedomorphosis – we also discovered the genomic mechanism by which the upregulation of the juvenile isoHb became canalized in Tibetan antelope. Specifically, we document how a reversible acclimatization response to acute hypoxia – as observed in modern-day sheep and goats – became genetically assimilated as an irreversible adaptation to chronic hypoxia.

## Results and Discussion

We characterized the genomic organization of globin genes in Tibetan antelope and other bovid artiodactyls using published genome assemblies (*17*). Among mammals, bovid artiodactyls are unusual in that the entire β-globin gene cluster has undergone multiple rounds of *en bloc* duplication involving the same set of pre- and post-natally expressed β-type globin genes (Fig. 1) (*18*-*20*). Cows (*Bos taurus*) possess two duplicated gene blocks, each containing separate paralogs of the β-globin gene, β^A^ and β^F^, in the 5’ and 3’ blocks, respectively (Fig. 1). As with other eutherian mammals, the product of β^A^ is incorporated into an adult-expressed isoHb, HbA (α_2_β^A^_2_), whereas the β^F^ gene has been recruited for prenatal expression, and is incorporated into a fetal isoHb, HbF (α_2_β^F^_2_) (*21*). Goats (*Capra hircus*) and sheep (*Ovis aries*) possess an additional gene block at the 5’ end of the cluster that contains a third β-globin paralog, β^C^ (Fig. 1) (*18*-*20*). Whereas the β^A^ and β^F^ genes in goats and sheep have retained the same developmental expression profiles as their respective orthologs in cow, the β^C^ gene has been recruited for a new ontogenetic stage of expression during the first few months of neonatal life, and its product is incorporated into a juvenile isoHb, HbC (α_2_β^C^_2_) (*22*).

**Fig. 1.**
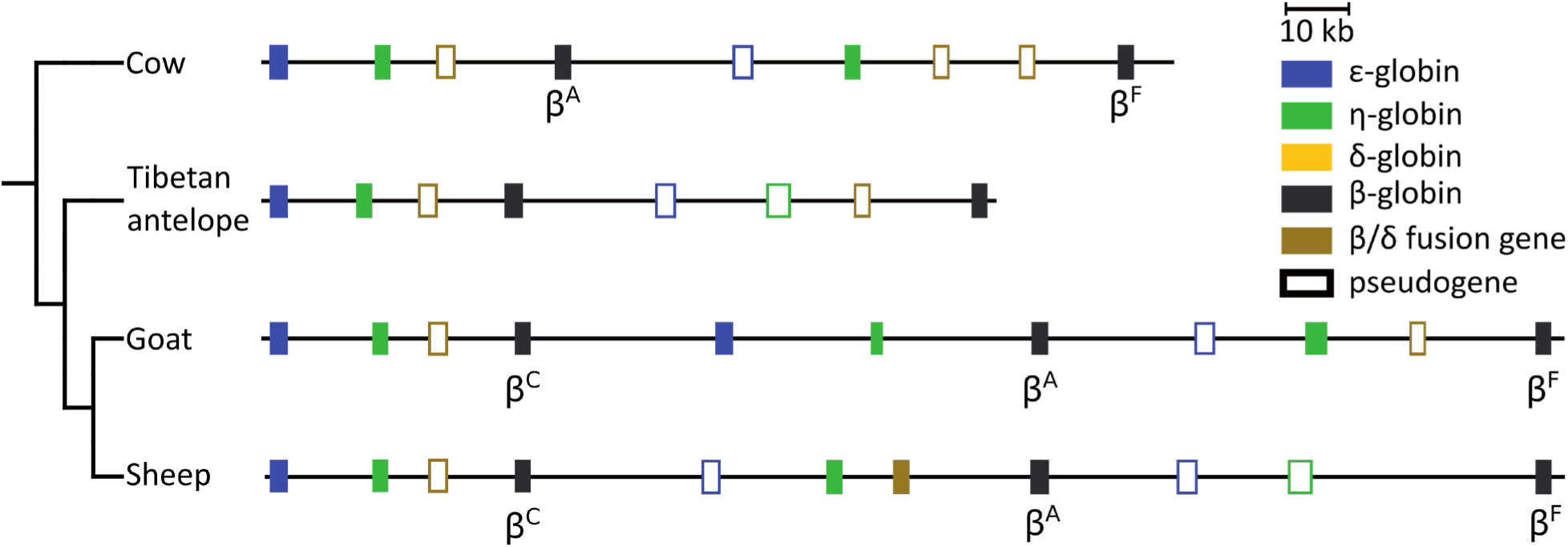
Genomic organization of bovid β-globin gene clusters. Colored boxes represent individual genes. Labels denote previously annotated β^C^-, β^A^-, and β^F^-globin genes.

The β-globin gene cluster of Tibetan antelope appears superficially similar to that of cow in terms of gene content (Fig. 1), suggesting that the Tibetan antelope inherited the same pair of β^A^– and β^F^–containing gene blocks. The alternative hypothesis is that Tibetan antelope inherited the additional *en bloc* duplication observed in goats and sheep, but one of the triplicated gene blocks was secondarily deleted, in which case the sole remaining pair of β-globin genes would be represented by one of three possible combinations: β^A^ + β^F^ (a reversion to the ancestral gene complement observed in cow), β^C^ + β^A^, or β^C^ + β^F^ (Fig. 2A-C). Either of the latter two combinations would implicate a novel isoHb profile that is not observed in other bovid taxa. To distinguish among these three alternative scenarios, we estimated the phylogeny of bovid β^C^, β^A^, and β^F^ genes and the pair of Tibetan antelope β-globin paralogs. Estimated phylogenies (Fig. 2D, supplementary Fig. S1) clearly demonstrate that the 5’ and 3’ β-globin genes of Tibetan antelope are orthologous to bovid β^C^ and β^F^, respectively, consistent with the scenario illustrated in Fig. 2B. This result indicates that Tibetan antelope inherited the triplicated set of β^C^-, β^A^-, and β^F^- containing gene blocks observed in goats and sheep (Fig. 1), and that the 5’ gene block containing β^A^ was secondarily deleted. This phylogenetic inference is unambiguously corroborated by patterns of conserved synteny and pairwise sequence matches (Fig. 3), as the β^C^- and β^F^-containing gene blocks of goat and sheep match the 5’ and 3’ gene blocks in Tibetan antelope. This comparative genomic analysis revealed that a ∼45 kb region of the Tibetan antelope β-globin gene cluster was deleted – a gene region that contained the ortholog of the β^A^ gene that encodes the β-chain of adult Hb in bovids and all other mammals.

**Fig. 2.**
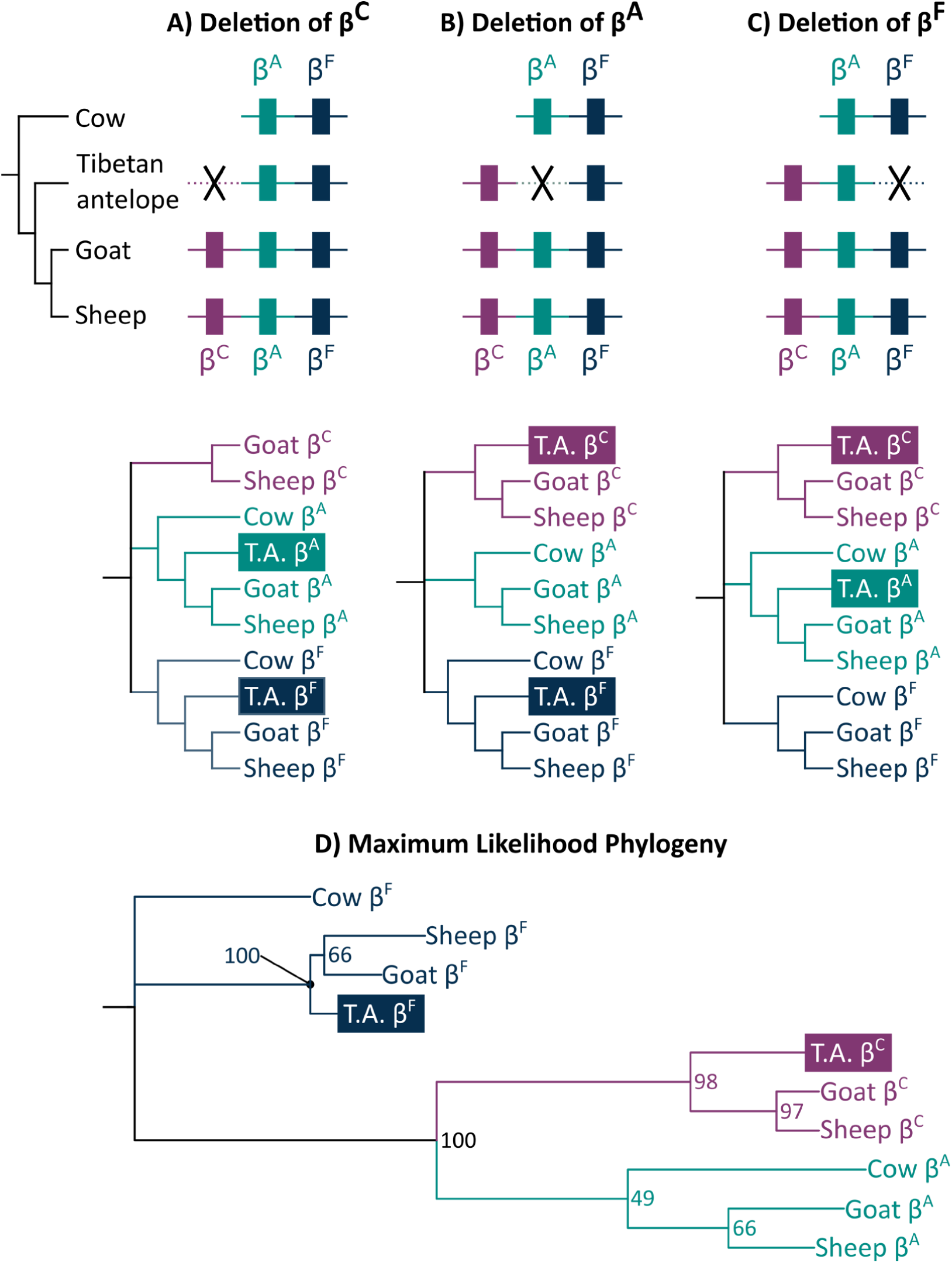
Phylogenetic analyses of bovid β^C^-, β^A^-, and β^F^-globin genes. Alternative histories of gene deletion in Tibetan antelope yield testable phylogenetic hypotheses: (A) deletion of β^C^, (B) deletion of β^A^, and (C) deletion of β^F^. (D) Estimated maximum likelihood phylogeny of bovid β-type globin genes indicates that Tibetan antelope has retained copies of β^C^ and β^F^, and that β^A^ has been secondarily lost. Bootstrap support values are shown for relevant nodes.

**Fig. 3.**
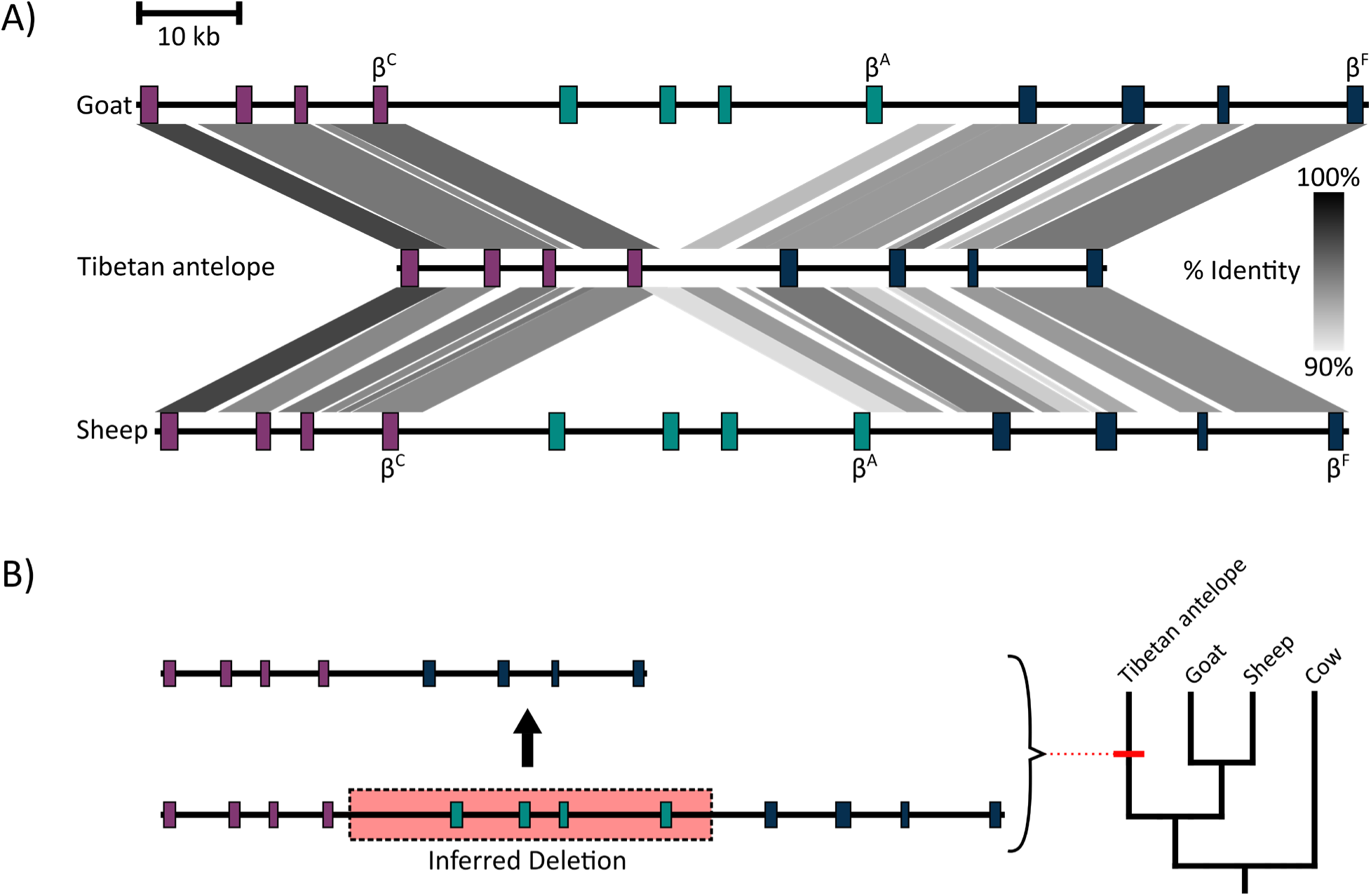
Analysis of pairwise sequence matches in chromosomal regions containing bovid β-globin gene clusters reveal a large-scale deletion in Tibetan antelope. Purple, green and blue colored boxes represent genes within the β^C^-, β^A^-, and β^F^-containing gene blocks, respectively. (A) Gray shading denotes percent sequence identity between homologous β-globin gene clusters. (B) A ∼45kb chromosomal deletion in the β-globin cluster of the Tibetan antelope lineage resulted in secondary loss of the β^A^-containing gene block.

Deletion of the adult β^A^ gene in the ancestor of Tibetan antelope effectively truncated the ancestral ontogeny of globin gene expression, such that juvenile HbC completely supplanted HbA in adult red blood cells. Thus, blood–O_2_ transport in Tibetan antelope has been juvenilized relative to the ancestral phenotype of adult bovids. To examine the effects of this paedomorphic change, we measured the oxygenation properties of purified recombinant Hb from Tibetan antelope and purified native Hbs from adult specimens of 10 other bovid species (Fig. 4; Table S1). The adult red cells of these other taxa contain HbA alone or in combination with HbC as a minor component. We measured the Hb–O_2_ affinity of purified total Hb from each bovid species in both the absence (stripped) and presence of 100 mM Cl^-^ (in the form of KCl). The stripped treatment provides a measure of intrinsic Hb–O_2_ affinity whereas the +KCl treatment provides a measure that is relevant to *in vivo* conditions in bovid red cells, as Cl^-^ ions are the principal allosteric regulators of Hb–O_2_ affinity (i.e., heme reactivity is modulated oxygenation-linked binding of Cl^-^ ions at sites remote from the heme iron)(*6, 21, 23*). Results of our *in vitro* experiments revealed that Hb of Tibetan antelope has a substantially higher O_2_–affinity than that of all other bovid taxa (Fig. 4; Table S1). Hbs of all taxa were similarly responsive to Cl^-^, as the average *P*_50_ (the *P*O_2_ yielding 50% Hb–O_2_ saturation) was 27.1% higher (i.e., Hb–O_2_ affinity was lower) in the +KCl treatment (Table S1).

**Fig. 4.**
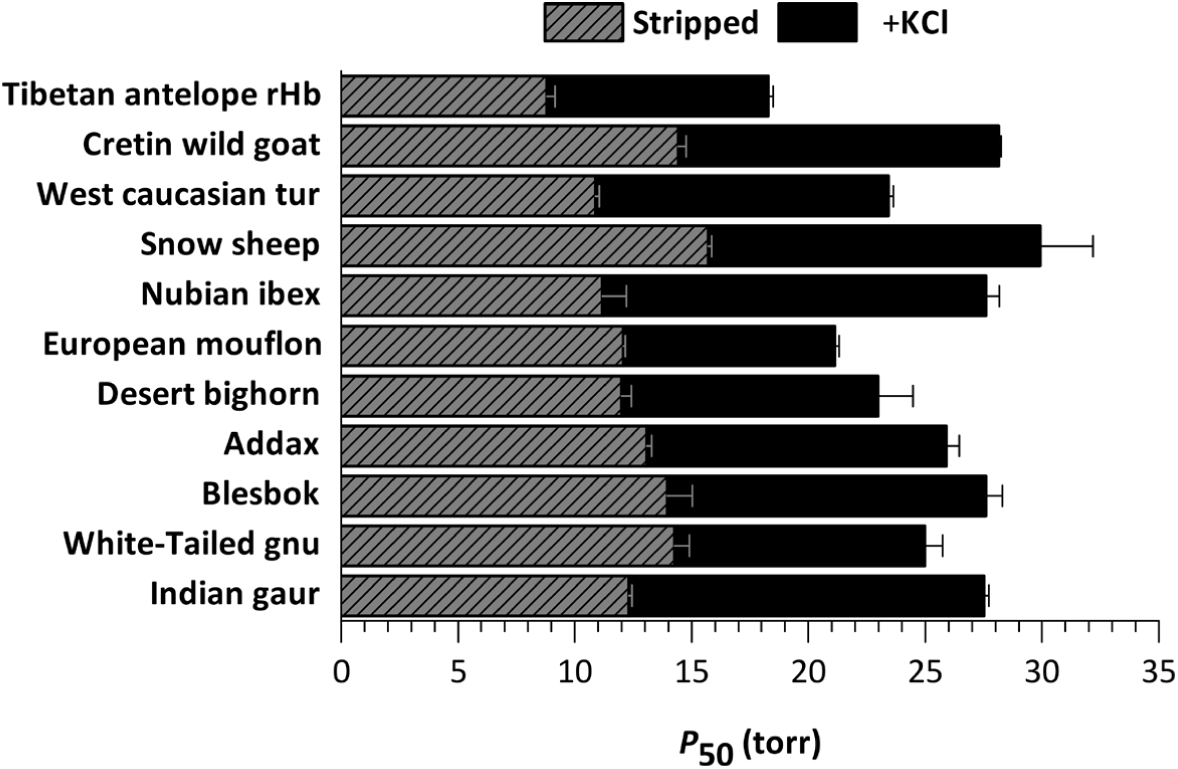
O_2_-affinities of total purified Hb from select bovid species. O_2_ tensions at half saturation (*P*_50_) for total Hb in the absence (stripped; gray hatched bars) and presence of 0.1 M KCl (black bars) at 37°C, pH 7.4 (0.1 mM Hb_4_). Values are shown as mean *P*_50_ ± s.e.m (n = 3).

As a follow-up experiment, we isolated and purified HbC and HbA from two of the bovid species expressing both components and we measured isoHb-specific O_2_–binding properties to determine how blood–O_2_ affinity would be affected by elimination of the major HbA isoHb (as would occur with the deletion of the β^A^ gene, leaving juvenile HbC as the sole-remaining isoHb). There was very little among-species variation in the measured O_2_–affinities of either juvenile HbC or adult HbA (Fig. S2; Table S1), but the O_2_–affinity of HbC exceeded that of HbA by a consistent margin (average 10.6 torr) in all species (Fig. S2; Table S1). Moreover, O_2_–affinity of HbC alone was always substantially higher than that of the composite HbA + HbC mixture (with the two isoHbs present in their naturally occurring relative abundance) (Fig. S2; Table S1), reflecting the fact that the lower affinity HbA is always present as the major isoHb in adult red cells (average HbA/HbC ratio = ∼80:20).

The higher Hb–O_2_ affinity of Tibetan antelope relative to that of other bovid species is entirely attributable to a difference in isoHb composition: they only express the high-affinity HbC instead of jointly expressing HbA and HbC (with the lower-affinity HbA present as the major isoHb). To infer the direction of evolutionary change in isoHb-specific O_2_–affinities, and to reconstruct the phenotypic effect of deleting β^A^-globin (thereby leaving HbC as the sole-expressed isoHb in adult red cells), we reconstructed the ancestral bovid β^A^ and β^C^ genes as well as their single-copy, pre-duplication progenitor (β^AC^) (Fig. 5; Fig. S3). Triangulated comparison of O_2_–affinities of the three recombinantly expressed ancestral isoHbs, AncHb-βA, AncHb-βC, and AncHb-β^AC^ (all of which had identical α-chains), revealed that the juvenile AncHb-β^C^ evolved a slight increase in O_2_–affinity relative to the estimated ancestral state (represented by AncHb-βAC), whereas adult HbA evolved a slight reduction in O_2_–affinity (Fig. 5). These data indicate that if HbA and HbC were present in a 80:20 ratio in the red cells of the Tibetan antelope ancestor (as in extant bovids), the deletion of β^A^-globin and the consequent elimination of HbA from the ‘HbA + HbC’ composite mixture would result in a 13.5% increase in Hb–O_2_ affinity in the presence of 100 mM Cl^-^ (*P*_50_ decreased from 18.5 to 16.0 torr).

**Fig. 5.**
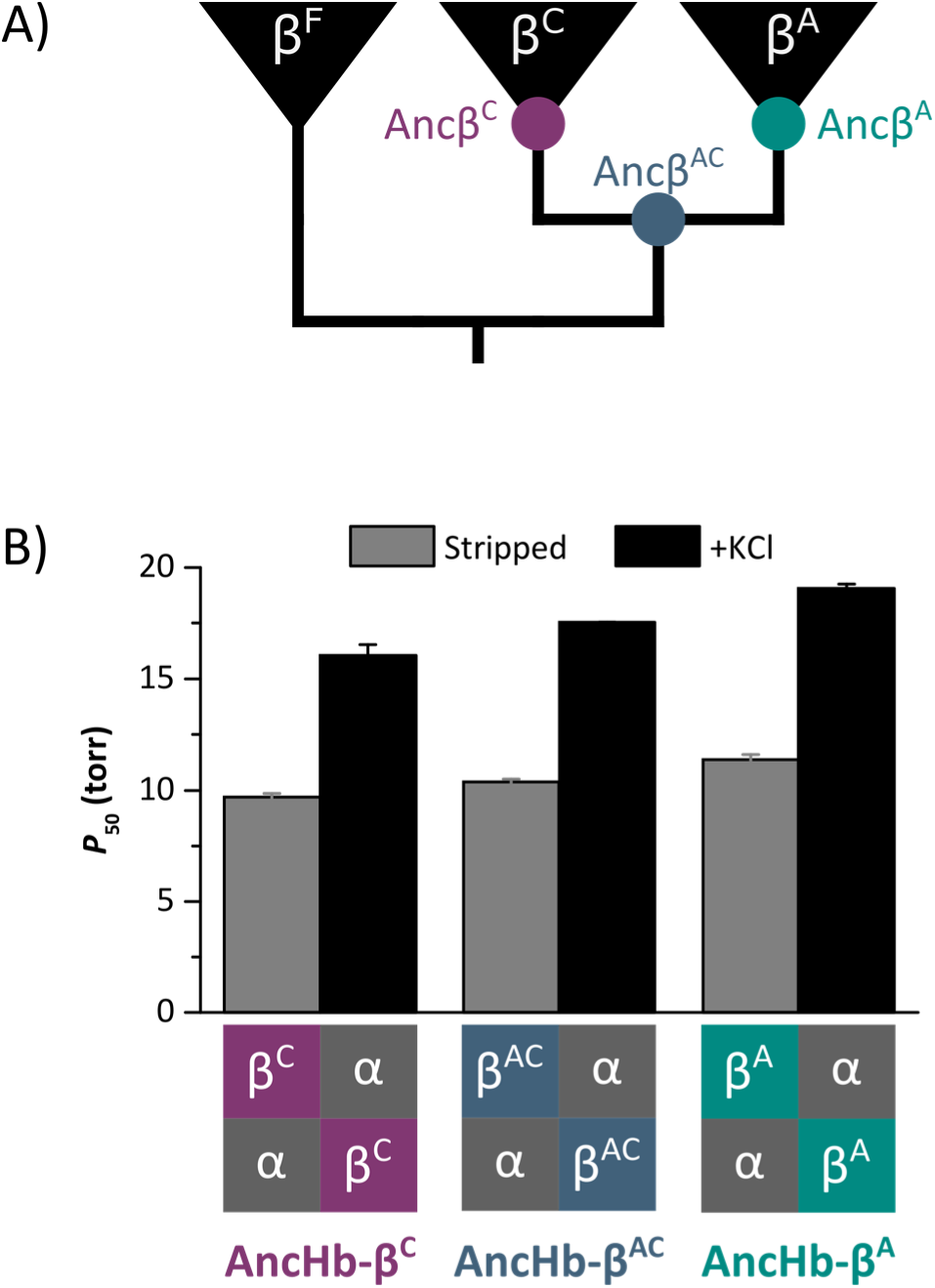
O_2_-affinities of reconstructed ancestral bovid isoHbs. (A) Reconstructed ancestral β-globin genes (β^C^, β^A^, and β^AC^) of bovids. (B) O_2_ tensions at half saturation (mean *P*_50_ ± s.e.m., n= 3) for recombinant ancestral isoHbs in the absence (stripped) and presence of 0.1 M KCl at 37°C, pH 7.4 (0.1 mM Hb_4_). Schematic diagrams show the subunit composition of the three ancestral isoHbs (which have identical α-chains and structurally distinct β-chains).

The derived blood phenotype of Tibetan antelope is consistent with the theoretical expectation that an increased Hb–O_2_ affinity is adaptive under conditions of severe hypoxia (especially in highly athletic species), and is consistent with patterns observed in other high-altitude mammals and birds that maintain especially high rates of aerobic metabolism (*24*-*32*). In other case studies of high-altitude vertebrates, evolved increases in Hb–O_2_ affinity have been traced to one or more amino acid substitutions in the α- and/or β-chain subunits of the α_2_β_2_ Hb tetramer (*6, 30-31*). Here we document a unique case in which an evolved change in Hb–O_2_ affinity has been accomplished via a heterochronic shift in globin gene expression, such that a high-affinity, juvenile isoHb supplants the lower-affinity, adult isoHb. This juvenilization of blood properties represents a novel mode of biochemical adaptation and highlights the utility of heterochrony as an adaptive mechanism, whereby “…the existing channel of ordinary ontogeny already holds the raw material in a particularly effective state for evolutionary change” (*2*).

There has been debate in the literature regarding the relative importance of regulatory vs. coding changes in genetic adaptation (*33*) and phenotypic evolution in general (*34*-*35*). In the case of Tibetan antelope, the evolved increase in Hb–O_2_ affinity was caused by an unusual combination of regulatory and structural changes. Specifically, a drastic regulatory switch in protein isoform expression (via truncation of the ancestral ontogeny of globin gene regulation) was caused by a large-scale chromosomal deletion, highlighting the surprising diversity of genetic mechanisms and substrates of phenotypic evolution.

## Materials and Methods

### Collection of blood samples

Frozen erythrocytes from 10 bovid species were provided by the San Diego Zoo Institute for Conservation Research (Uniform Biological Material Transfer Agreement BR2017063). This included six species in the subfamily Caprinae (*Capra aegagrus cretica, Capra caucasica caucasica, Ovis nivicola, Capra nubiana, Ovis orientalis musimon* and *Ovis canadensis nelsoni*), two species in the subfamily Alcelaphinae (*Damaliscus pygargus phillipsi* and *Connochaetes gnou*), and one species of each from Hippotraginae (*Addax nasomaculatus*) and Bovinae (*Bos gaurus*)

### Sequencing of bovid globin genes

RNA was extracted from ∼100 ul of flash frozen erythrocytes using an RNeasy Universal Plus Mini Kit (Qiagen). cDNA was synthesized from freshly prepared RNA using Superscript IV Reverse transcriptase (Invitrogen). Gene specific primers were used to amplify the a- and b-type globin transcripts. PCR reactions were conducted using 1 ml of cDNA template in 0.2 ml tubes containing 25 µl of reaction mixture (0.5 µl of each dNTP (2.5 mM), 2.5 µl of 10x Reaction Buffer (Invitrogen), 0.75 µl of 50 mM MgCl_2_, 1.25 µl of each primer (10 pmol/µl), 1 µl of Taq polymerase (Invitrogen) and 16.75 µl of ddH_2_O), using an Eppendorf Mastercycler^®^ Gradient thermocycler. Following a 5-min denaturation period at 94°C, the desired products were amplified using a cycling profile of 94°C for 30 sec; 53-65°C for 30 sec; 72°C for 45 sec for 30 cycles followed by a final extension period of 5 min at 72°C. Amplified products were run on a 1.5% agarose gel and bands of the correct size were subsequently excised and purified using Zymoclean Gel DNA recovery columns (Zymo Research). Gel-purified PCR products were ligated into pCR™4-TOPO® vectors using a TOPO™ TA Cloning™ Kit and were then transformed into One Shot™ TOP10 Chemically Competent *E. coli* (Thermo Fisher Scientific). Three to six transformed colonies were cultured in 5 ml of LB medium and plasmids were subsequently purified with a GeneJET Plasmid Miniprep kit (Thermo Fisher Scientific). Purified plasmids were sequenced by Eurofins Genomics (Fig. S3).

### Sequence analyses

Genomic sequences containing the complete α- and β-globin gene clusters for the domestic goat (*Capra hircus*), sheep (*Ovis aries*), cow (*Bos taurus*), and Tibetan antelope (*Panthelops hodgsonii*) were mined from GenBank (Table S2). Sequence identity between bovid chromosomal regions containing the β-globin gene clusters was calculated using Blastn and conserved synteny was visualized using Easyfig 2.1 (*36*). Coding sequences of α- and β-globin genes were extracted from genomic and cDNA sequences mined from GenBank (Table S2) and combined with cDNA sequences collected above (Fig. S3). These sequences were aligned using MUSCLE (*37*) and were used for phylogenetic tree construction. The best fitting codon substitution model and initial tree search were estimated using IQ-TREE with the options -st CODON, -m TESTNEW, -allnni, and -bnni (*38, 39*). Initial trees were then subjected to 1000 ultrafast bootstrap replicates (*40*). Bootstrap consensus trees (Figs. S1A, S1B) were used to estimate ancestral globin sequences using IQ-TREE with the option -asr (Figs. S1C, S3). As bovid β^C^-globins are truncated by 9bp (relative to β^A^), the ancestral reconstruction of indels in the β-globin gene tree was performed by FastML (*41*).

### Protein purification

Blood samples (∼200μl) were added to a 5x volume of ice-cold water and incubated on ice for 30 minutes to lyse the red blood cells. Samples were centrifuged at 20,000 x g for 10 minutes to remove cell debris. Buffer was added to the supernatants to a final concentration of 0.01 M HEPES/0.2 M NaCl (pH 7.4) and passed through a PD-10 desalting column (GE Healthcare) equilibrated with 25 ml of 0.01 M HEPES/0.5mM EDTA (pH 7.2) to remove intracellular cofactors. Desalted lysates were loaded onto a HiTrap SP cation exchange column (GE Healthcare) and isoHbs were eluted using a linear pH gradient (0.01 M HEPES/0.5mM EDTA, pH 7.2 – 7.7). For each species, a subsample of each isoHb was pooled to create a “Total Hb” solution. Each Hb solution was then desalted using a PD-10 column (GE Healthcare) equilibrated with 0.01 M HEPES/0.5mM EDTA (pH 7.4) and eluates were concentrated using Amicon Ultra-4 Centrifugal Filter Units (Millipore).

### Measuring O_2_–binding properties of purified Hbs

O_2_–equillibrium curves for purified Hb solutions (0.1 mM hemoglobin in 0.1 M HEPES/0.05 M EDTA buffer, pH 7.4) were measured at 37**°**C using a Blood Oxygen Binding System (Loligo Systems). O_2_–equillibrium curves were measured in the absence (stripped) and presence of chloride ions (0.1 M KCl). Each Hb solution was sequentially equilibrated with three to five different O_2_ tensions (*P*O_2_) at saturation levels between 30 to 70% while the absorbance was continually monitored at 430 nm (deoxy peak) and 421 nm (oxy/deoxy isobestic point)(*42-44*). Hill plots (log[fractional saturation/[1-fractional saturation]] vs log*P*O_2_) constructed from these measurements were used to estimate the *P*O_2_ at half saturation (*P*_50_) and the cooperativity coefficient (*n*_50_) from the χ-intercept and slope of these plots, respectively. O_2_–equilibrium curves for each Hb solution were measured in triplicate and *P*_50_ is reported as mean ± s.e.m. (Table S1).

### Construction of Hb expression vector

Globin sequences for domestic goat, Tibetan antelope, as well as the reconstructed ancestral globins were synthesized by GeneArt Gene Synthesis (Thermo Fisher Scientific) after optimizing the nucleotide sequences in accordance with *E. coli* codon preferences. The synthesized globin gene cassette was cloned into a custom pGM vector system along with the *methionine aminopeptidase* (MAP) gene, as described previously (*45*).

### Expression and purification of recombinant Hbs

Recombinant Hb expression was carried out in the *E. coli* JM109 (DE3) strain as described previously (*45-47*). Bacterial cell lysates were then loaded onto a HiTrap Q HP anion exchange column (GE Healthcare) and were then equilibrated with 20 mM Tris/0.5 mM EDTA (pH 8.3) and eluted with a linear gradient of 0 - 0.25 M NaCl. Hb-containing fractions were then loaded on to a HiTrap SP HP cation exchange column (GE Healthcare) and eluted with a linear pH gradient (pH 6.8 to 8.4). Eluted Hb factions were concentrated using Amicon Ultra-4 Centrifugal Filter Units (Millipore) and oxygenation properties were measured as described above.

## Acknowledgments

We thank C. Natarajan for assistance in the laboratory and K. Campbell for comments on this manuscript.

## Funding

This study was funded by grants to JFS from the National Institutes of Health (HL087216) and the National Science Foundation (MCB-1517636 and OIA-1736249).

## Author contributions

AVS and JFS designed the study, AVS acquired the data, AVS and JFS analyzed the data and wrote the manuscript.

## Competing interests

Authors declare no competing interests.

## Data and materials availability

Sequence data collected as part of this study are deposited to GenBank under accession numbers XXXXXXX-YYYYYYY.

## Supplementary Materials

Figures S1-S3

Tables S1-S2

## References

1. S. J. Gould, Ontogeny and phylogeny. (Harvard University Press, Cambridge, MA, (1977).

2. S. J. Gould, The structure of evolutionary theory (Harvard University Press, Cambridge, MA, 2002).

3. R. E. Weber, Hemoglobin-based O2 transfer in viviparous animals. Israel J. Zool. (1994).

4. T. Brittain, Molecular aspects of embryonic hemoglobin function. Mol. Aspects. Med. 23, 293–342 (2002).

5. J. F. Storz, Gene duplication and evolutionary innovations in hemoglobin-oxygen transport. Physiol. 31, 223–232 (2016).

6. J. F. Storz, Hemoglobin: Insights into Protein Structure, Function, and Evolution (Oxford University Press, USA, 2019).

7. D. S. Vinjamur, D. E. Bauer, S. H. Orkin, Recent progress in understanding and manipulating haemoglobin switching for the haemoglobinopathies. Br. J. Haematol. 180, 630–643 (2018).

8. J. F. Storz, G. R. Scott, Z. A. Cheviron. 2010. Phenotypic plasticity and genetic adaptation to high-altitude hypoxia in vertebrates. J. Exp. Biol. 213, 4125–4136 (2010).

9. J. F. Storz, Hemoglobin-oxygen affinity in high-altitude vertebrates: is there evidence for an adaptive trend? J. Exp. Biol. 219, 3190–3203 (2016).

10. J. F. Storz, G. R. Scott. Life ascending: mechanism and process in physiological adaptation to high-altitude hypoxia. Annu. Rev. Ecol. Evol. Syst. 50, 503–526 (2019).

11. P. Dominelli, C. C. Wiggins, S. E. Baker, J. R. A. Shepherd, S. K. Roberts, T. K. Roy, T. B. Curry, J. D. Hoyer, J. L. Oliveira, M. J. Joyner. Influence of high affinity hemoglobin on the response to normoxic and hypoxic exercise. J. Physiol. https://doi.org/10.1113/JP279161 (2020).

12. M. H. Blunt, T. H. Huisman, J. P. Lewis, The production of haemoglobin C in adult sheep and goats. Aust. J. Exp. Biol. Med. Sci. 47, 601–611 (1969).

13. M. H. Blunt, M. Perry, R. Lane, The production of haemoglobin C by sheep at simulated high altitude. R. Vet. Sci. 11, 191–193 (1970).

14. A. L. Cymerman. J. T. Reeves, J. R. Sutton, P. B. Rock, B. M. Groves, M. K. Malconian, P. M. Young, P. D. Wagner, C. S. Houston. Operation Everest II: maximal oxygen uptake at extreme altitude. J. Appl. Physiol. 66, 2446–2453 (1989).

15. M.A. Liu, S. Mahalingam, P. Patel, A. D. Connaty, C. M. Ivy, Z. A. Cheviron, J. F. Storz, G. B. McClelland, G. R. Scott. High-altitude ancestry and hypoxia acclimation have distinct effects on exercise capacity and muscle phenotype in deer mice. Am. J. Physiol. Regul. Integr. Comp. Physiol. 308, R779–R791 (2015).

16. H. B. Zhuang, W. D. Li, Z. H. Liu, Tibetan antelope (*Pantholops hodgsoni*). Chin. J. Zool. 38, 74 (2003)

17. R. L. Ge, Q. Cai, Y. Y. Shen, A. San, L. Ma, Y. Zhang, X. Yi, Y. Chen, L. Yang, Y. Huang, R. He, Y. Hui, M. Hao, Y. Li, B. Wang, X. Ou, J. Xu, Y. Zhang, K. Wu, C. Geng, W. Zhou, T. Zhou, D. M. Irwin, Y. Yang, L. Ying, H. Bao, J. Kim, D. M. Larkin, J. Ma, H. A. Lewin, J. Xing, R. N. Platt 2nd, D. A. Ray, L. Auvil, B. Capitanu, X. Zhang, G. Zhang, R. W. Murphy, J. Wang, Y. P. Zhang, J. Wang, Draft genome sequence of the Tibetan antelope. Nat. Com. 4, 1–7 (2013).

18. T. M. Townes, M. C. Fitzgerald, J. B. Lingrel, Triplication of a four-gene set during evolution of the goat β-globin locus produced three genes now expressed differentially during development. Proc. Nat. Acad. Sci. 81, 6589 (1984).

19. J. C. Schimenti, C. H. Duncan, Structure and organization of the bovine β-globin genes. Mol. Biol. Evol. 2, 514–525 (1985).

20. M. J. Gaudry, J. F. Storz, G. T. Butts, K. L. Campbell, F. G. Hoffmann, Repeated evolution of chimeric fusion genes in the β-globin gene family of laurasiatherian mammals. Gen. Biol. Evol. 6, 1219–1234 (2014).

21. M. E. Clementi, R. Scatena, A. Mordente, Oxygen transport by fetal bovine hemoglobin. J. Mol. Biol. 255, 229–234 (1996).

22. M. H. Blunt, T. H. J. Huisman, “The Haemoglobins of Sheep” in The Blood of Sheep: Composition and function (Springer, New York, 1975).

23. H. F. Bunn, Differences in the interaction of 2,3-diphosphoglycerate with certain mammalian hemoglobins. Science 172, 1049–1050 (1971).

24. J. F. Storz, A. M. Runck, H. Moriyama, R. E. Weber, A. Fago, Genetic differences in hemoglobin function between highland and lowland deer mice. J. Exp. Biol. 213, 2565–2574 (2010).

25. J. Projecto-Garcia, C. Natarajan, H. Moriyama, R. E. Weber, A. Fago, Z. A. Cheviron, R. Dudley, J. A. McGuire, C. C. Witt, J. F. Storz. Repeated elevational transitions in hemoglobin function during the evolution of Andean hummingbirds. Proc. Natl. Acad. Sci. USA 110, 20669–20674 (2013).

26. S. C. Galen, C. Natarajan, H. Moriyama, R. E. Weber, A. Fago, P. M. Benham, A. N. Chavez, Z. A. Cheviron, J. F. Storz. Contribution of a mutational hot spot to hemoglobin adaptation in high-altitude Andean house wrens. Proc. Natl. Acad. Sci. USA 112, 13958–13963 (2015).

27. C. Natarajan, J. Projecto-Garcia, H. Moriyama, R. E. Weber, V. Muñoz-Fuentes, A. J. Green, C. Kopuchian, P. L. Tubaro, L. Alza, M. Bulgarella, M. M. Smith, R. E. Wilson, A. Fago, K. G. McCracken, J. F. Storz, Convergent evolution of hemoglobin function in high-altitude Andean waterfowl involves limited parallelism at the molecular sequence level. PLoS Genet. 11, e1005681 (2015).

28. C. Natarajan, F. G. Hoffmann, H. C. Lanier, C. J. Wolf, Z. A. Cheviron, M. L. Spangler, R. E. Weber, A. Fago, J. F. Storz. Intraspecific polymorphism, interspecific divergence, and the origins of function-altering mutations in deer mouse hemoglobin. Mol. Biol. Evol. 32, 978–997 (2015).

29. D. M. Tufts, C. Natarajan, I. G. Revsbech, J. Projecto-Garcia, F. G. Hoffmann, R. E. Weber, A. Fago, H. Moriyama, J. F. Storz. Epistasis constrains mutational pathways of hemoglobin adaptation in high-altitude pikas. Mol. Biol. Evol. 32, 287–298 (2015).

30. C. Natarajan, F. G. Hoffmann, R. E. Weber, A. Fago, C. C. Witt, J. F. Storz. Predictable convergence in hemoglobin function has unpredictable molecular underpinnings. Science 354, 336–339 (2016).

31. X. Zhu, Y. Guan, A. V. Signore, C. Natarajan, S. G. DuBay, Y. Cheng, N. Han, G. Song, Y. Qu, H. Moriyama, F. G. Hoffmann, A. Fago, F. -M. Lei, J. F. Storz. Divergent and parallel routes of biochemical adaptation in high-altitude passerine birds from the Qinghai-Tibet Plateau. Proc. Natl. Acad. Sci. USA 115, 1865–1870 (2018).

32. A. V. Signore, Y. -Z. Yang, Q. -Y. Yang, G. Qin, H. Moriyama, R. -L. Ge, J. F. Storz. Adaptive changes in hemoglobin function in high-altitude Tibetan canids were derived via gene conversion and introgression. Mol. Biol. Evol. 36, 2227–2237 (2019).

33. H. E. Hoekstra, J. A. Coyne, The locus of evolution: evo devo and the genetics of adaptation. Evolution 61, 995–1016 (2007).

34. S. B. Carroll, Evo-devo and an expanding evolutionary synthesis: a genetic theory of morphological evolution. Cell 134, 25–36 (2008).

35. D. L. Stern, V. Orgogozo, Is genetic evolution predictable? Science 323, 746–751 (2009).

36. M. J. Sullivan, N. K. Petty, S. A. Beatson, Easyfig: a genome comparison visualizer. Bioinformatics. 27, 1009–1010 (2011).

37. R. C. Edgar, MUSCLE: multiple sequence alignment with high accuracy and high throughput. Nuc. Acid. Res. 32, 1792–1797 (2004).

38. L.-T. Nguyen, H. A. Schmidt, A. von Haeseler, B. Q. Minh, IQ-TREE: A fast and effective stochastic algorithm for estimating maximum-likelihood phylogenies. Mol. Biol. Evol. 32, 268–274 (2015).

39. D. T. Hoang, O. Chernomor, UFBoot2: Improving the ultrafast bootstrap approximation. Mol. Biol. Evol., 35, 518–522 (2018).

40. S. Kalyaanamoorthy, B. Q. Minh, T. K. F. Wong, A. von Haeseler, L. S. Jermiin, ModelFinder: fast model selection for accurate phylogenetic estimates. Nat. Met. 14, 587–589 (2017).

41. H. Ashkenazy, O. Penn, A. Doron-Faigenboim, O. Cohen, G. Cannarozzi, O. Zomer, T. Pupko, FastML: a web server for probabilistic reconstruction of ancestral sequences. Nuc. Acid. Res. 40, W580–W584 (2012).

42. M. T. Grispo, C. Natarajan, J. Projecto-Garcia, H. Moriyama, R. E. Weber, J. F. Storz, Gene duplication and the evolution of hemoglobin isoform differentiation in birds. J. Biol. Chem. 287, 37647–37658 (2012).

43. J. F. Storz, C. Natarajan, H. Moriyama, F. G. Hoffmann, T. Wang, A. Fago, H. Malte, J. Overgaard, R. E. Weber, Oxygenation properties and isoform diversity of snake hemoglobins. Am. J. Physiol. Regul. Integr. Comp. Physiol. 309, R1178–R1191 (2015).

44. A. Fago, C. Natarajan, M. Pettinati, F. G. Hoffmann, T. Wang, R. E. Weber, S. I. Drusin, F. Issoglio, M. A. Martí, D. A. Estrin, J. F. Storz, Structure and function of crocodilian hemoglobins and allosteric regulation by chloride, ATP, and CO2. Am. J. Physiol. Regul. Integr. Comp. Physiol. 318, R657–R667 (2020).

45. C. Natarajan, X. Jiang, A. Fago, R. E. Weber, H. Moriyama, J. F. Storz, Expression and purification of recombinant hemoglobin in *Escherichia coli*. PLoS One. 6, e20176 (2011).

46. C. Natarajan, N. Inoguchi, R. E. Weber, A. Fago, H. Moriyama, J. F. Storz, Epistasis among adaptive mutations in deer mouse hemoglobin. Science 340, 1324–1327 (2013).

47. Z. A. Cheviron, C. Natarajan, J. Projecto-Garcia, D. K. Eddy, J. Jones, M. D. Carling, C. C. Witt, H. Moriyama, R. E. Weber, A. Fago, J. F. Storz, Integrating evolutionary and functional tests of adaptive hypotheses: a case study of altitudinal differentiation in hemoglobin function in an Andean sparrow, *Zonotrichia capensis*. Mol. Biol. Evol. 31, 2948–2962 (2014).

