## Supplementary Materials for "Biochemical paedomorphosis and genetic assimilation in the hypoxia adaptation of Tibetan antelope"

**This PDF file includes:**

Figs. S1 to S3
Tables S1 to S2

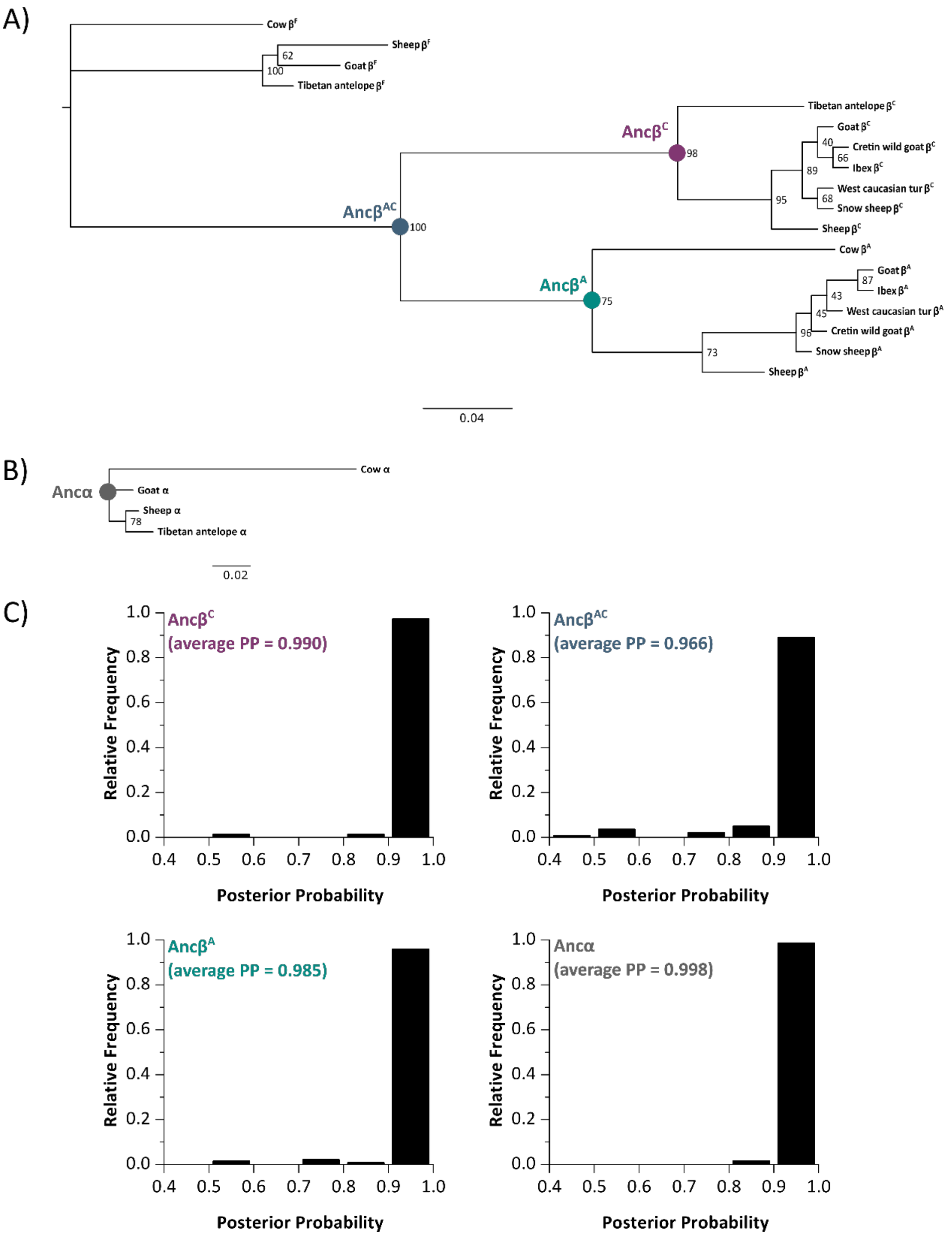


**Fig. S1.** Ancestral sequence reconstructions of bovid globin genes. Estimated maximum likelihood phylogenies of bovid (A) β-globin and (B) α-globin genes. Filled circles represent nodes for which ancestral sequences were reconstructed. Scale bars denote the mean number of substitutions per site. (C) Relative frequencies of posterior probabilities for each reconstructed codon. Histogram label colors correspond to node colors in A and B.

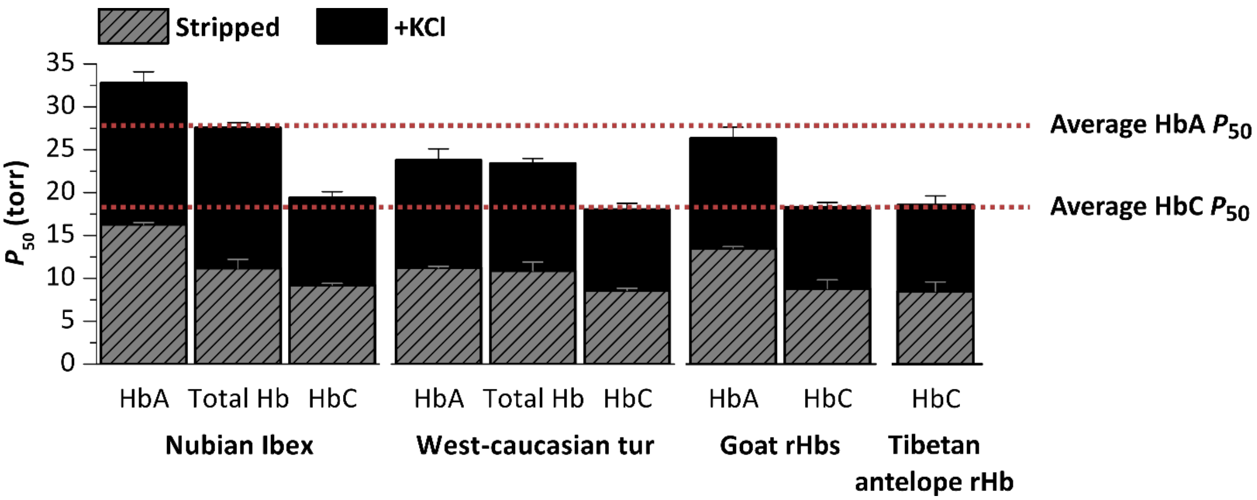


**Fig. S2.** $O_2$–affinities of purified HbA, HbC, and total Hb (HbA + HbC) from select bovid species. $O_2$ tensions at half saturation ($P_{50}$, torr; mean ± s.e.m., n=3) are shown for individual isoHbs and total Hb in the absence and presence of 0.1 M KCl at 37°C, pH 7.4 (0.1 mM $Hb_4$).

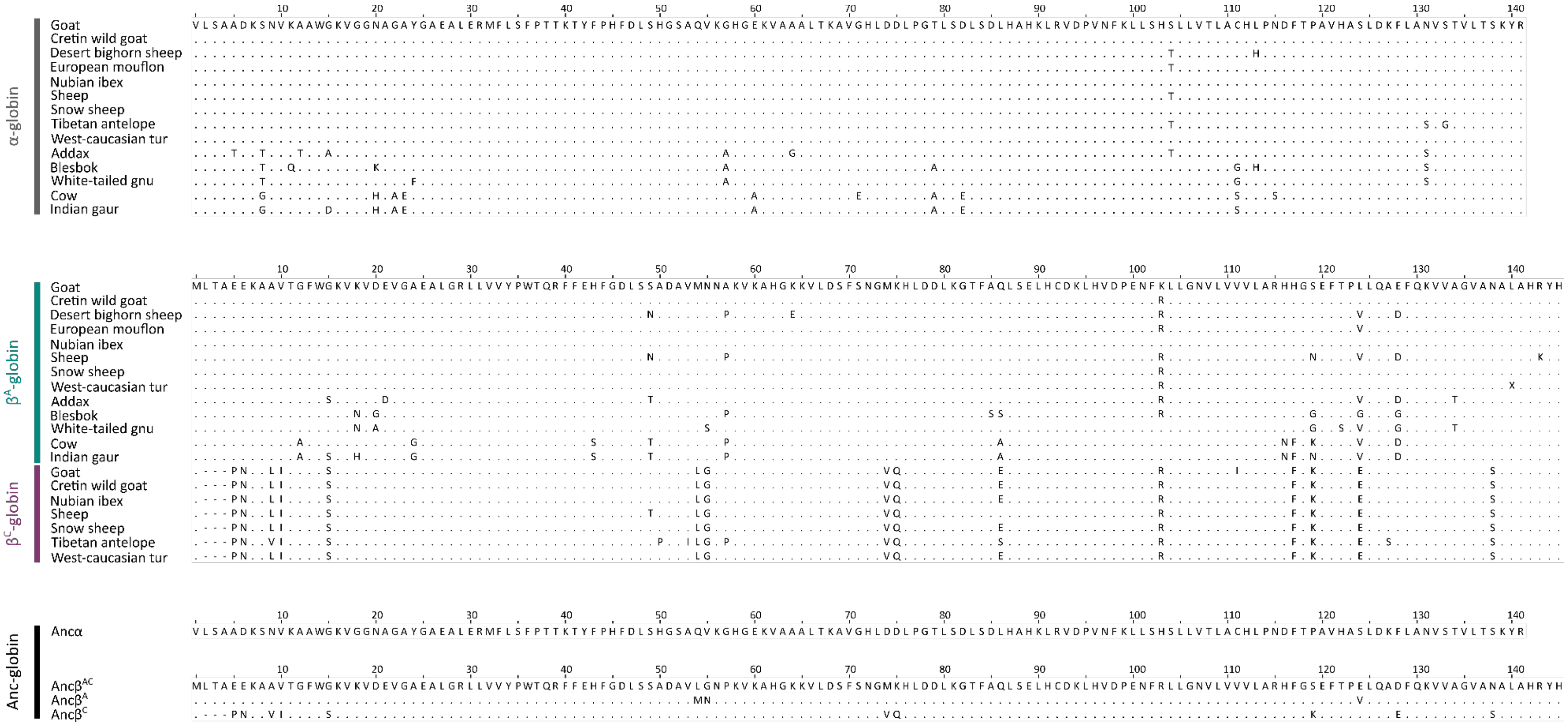


**Fig. S3.** Amino acid variation in postnatally expressed α- and β-type globins of select bovid species.

**Table S1.** Oxygenation properties of purified bovid hemoglobins. $P_{50}$ values are reported as mean ± s.e.m.

| **Hemoglobin solution** | **Species** | **$P_{50}$, Stripped (torr)** | | | **$P_{50}$, +KCl (torr)** | | | **KCl Effect ($\log P_{50}$[+KCl] - $\log P_{50}$[Stripped])** |
|---|---|---|---|---|---|---|---|---|
| Total Hb | European mouflon, *Ovis orientalis musimon* | 12.08 | ± | 0.09 | 21.12 | ± | 0.20 | 0.24 |
| | Snow sheep, *Ovis nivicola* | 15.72 | ± | 0.12 | 29.91 | ± | 2.28 | 0.28 |
| | Desert bighorn sheep, *Ovis canadensis nelsoni* | 12.00 | ± | 0.42 | 22.98 | ± | 1.50 | 0.28 |
| | Cretin wild goat, *Capra aegagrus cretica* | 14.43 | ± | 0.33 | 28.14 | ± | 0.10 | 0.29 |
| | Nubian ibex, *Capra nubiana* | 11.20 | ± | 1.01 | 27.59 | ± | 0.57 | 0.39 |
| | West caucasian tur, *Capra caucasica* | 10.90 | ± | 0.15 | 23.41 | ± | 0.21 | 0.33 |
| | White-tailed gnu, *Connochaetes gnou* | 14.27 | ± | 0.64 | 24.97 | ± | 0.78 | 0.24 |
| | Blesbok, *Damaliscus pygargus phillipsi* | 13.96 | ± | 1.09 | 27.59 | ± | 0.71 | 0.30 |
| | Addax, *Addax nasomaculatus* | 13.07 | ± | 0.22 | 25.89 | ± | 0.62 | 0.30 |
| | Indian gaur, *Bos gaurus* | 12.32 | ± | 0.14 | 27.50 | ± | 0.24 | 0.35 |
| | Tibetan antelope rHb, *Panthalops hodgsonii* | 8.53 | ± | 1.10 | 18.59 | ± | 1.07 | 0.34 |
| Pure HbA | Nubian ibex, *Capra nubiana* | 16.30 | ± | 0.19 | 32.79 | ± | 1.30 | 0.30 |
| | West caucasian tur, *Capra caucasica* | 11.22 | ± | 0.42 | 23.79 | ± | 1.12 | 0.33 |
| | Goat rHb, *Capra hircus* | 13.51 | ± | 0.21 | 26.33 | ± | 0.53 | 0.29 |
| Pure HbC | Nubian ibex, *Capra nubiana* | 9.21 | ± | 0.22 | 19.41 | ± | 0.71 | 0.32 |
| | West caucasian tur, *Capra caucasica* | 8.62 | ± | 0.12 | 18.03 | ± | 0.57 | 0.32 |
| | Goat rHb, *Capra hircus* | 8.82 | ± | 0.33 | 18.29 | ± | 0.21 | 0.32 |

**Table S2.** Genomic sequences of the α- and β-globin gene clusters of bovid species that were included in the phylogenetic analyses.

| **Species** | **α-globin accession #s** | **β-globin accession #s** |
|---|---|---|
| Cow, *Bos taurus* | NC_037352.1 | NC_037342.1 |
| Goat, *Capra hircus* | NC_030832.1 | NC_030822.1 |
| Sheep, *Ovis aeries* | NC_040275.1 | NC_040266.1 |
| Tibetan antelope, *Panthalops hodgsonii* | JF811751.1 | JX276960.1 |
| | DQ650713.1 | HQ897270.1 |
| | NW_005806747 | AGTT01081104.1 |
| | | AGTT01081105.1 |
| | | AGTT01081106.1 |
| | | AGTT01081107.1 |
| | | AGTT01081108.1 |
| | | AGTT01081109.1 |
| | | AGTT01081110.1 |
| | | AGTT01081111.1 |
